# FOXO1 promotes HIV Latency by suppressing ER stress in T cells

**DOI:** 10.1101/2020.04.23.058123

**Authors:** Albert Vallejo-Gracia, Irene P. Chen, Rosalba Perrone, Emilie Besnard, Daniela Boehm, Emilie Battivelli, Tugsan Tezil, Karsten Krey, Kyle A. Raymond, Philip A. Hull, Marius Walter, Ireneusz Habrylo, Andrew Cruz, Steven Deeks, Satish Pillai, Eric Verdin, Melanie Ott

**Author notes:** Contributed equally (Irene P. Chen, Rosalba Perrone). Correspondence to Melanie Ott.

## Abstract

Quiescence is a hallmark of CD4^+^ T cells latently infected with HIV-1. While reversing this quiescence is an effective approach to reactivate latent HIV from T cells in culture, it can cause deleterious cytokine dysregulation in patients. Here we report that FOXO1, a key regulator of T-cell quiescence, promotes latency and suppresses productive HIV infection. In resting T cells, FOXO1 inhibition induces ER stress and activates two associated transcription factors: activating transcription factor 4 (ATF4) and nuclear factor of activated T cells (NFAT). Both factors associate with HIV chromatin and are necessary for HIV reactivation. Indeed, inhibition of PKR-like endoplasmic reticulum kinase (PERK), a known link between ER stress and ATF4, and calcineurin, a calcium-dependent regulator of NFAT, synergistically suppress HIV reactivation induced by FOXO1 inhibition. Thus, our studies uncover a novel link between FOXO1, ER stress, and HIV infection that could be therapeutically exploited to selectively reverse T-cell quiescence and reduce the size of the latent viral reservoir.

## Main

The major barrier to eradicating human immunodeficiency virus (HIV-1) from infected patients is the persistence of latently infected cells, primarily in memory CD4^+^ T cells. These cells are rare, long-lived, and generally quiescent. Although anti-retroviral therapy (ART) effectively suppresses HIV replication, therapy interruption results in reactivation of the latent reservoir, necessitating life-long treatment, and there is no cure^1,2^. One strategy to eliminate latently infected cells is to activate virus production using latency reversing agents (LRAs) with the objective of triggering cell death through virus-induced cytolysis or immune clearance. Efficient viral reactivation requires a calibrated degree of activation that avoids inducing polyclonal T cell activation or a cytokine storm that would negate the potential benefit. Known LRAs target various pathways to induce viral reactivation, and include PKC agonists (e.g. Prostratin), epigenetic modulators (e.g. HDAC or bromodomain inhibitors), and modulators of the PI3K or mTOR signaling pathways^3–6^. However, to date, clinical trials of existing LRAs have not demonstrated successful reduction of the latent reservoir^7^. Thus, a better understanding of how HIV latency is established and maintained is needed to improve the design of interventions.

The Forkhead box O (FOXO) protein family comprises evolutionarily conserved transcription factors^8^. FOXO-mediated gene regulation is determined in part by localization; upon phosphorylation by upstream kinases, they exit the nucleus and remain inactive. Different human FOXO isoforms influence multiple pathways, including cell cycle arrest, glucose metabolism, oxidative stress regulation, apoptosis, and the DNA damage response^9–11^. In the immune system, FOXO proteins regulate a set of genes involved in maintaining quiescence and cell-fate differentiation in CD4^+^ and CD8^+^ T cells^12–14^. Furthermore, inhibition of FOXO1 in naïve and memory CD8^+^ T cells promotes a more effector-like and cytotoxic phenotype in the context of immune aging and chronic infection^15^.

FOXO3 is downregulated in memory CD4^+^ T cells of elite controller HIV patients, promoting the persistence of these cells^16^. Furthermore, FOXO proteins negatively regulate HIV transcription through Tat-mediated repression^17^, and FOXO1 inhibition accelerates productive infection^18^. Because FOXO activity, especially FOXO1, promotes and maintains the quiescent state of memory CD4^+^ T cells, in which HIV establishes latency, we hypothesized that FOXO1, directly or indirectly, induces HIV latency. Here we identify a pathway that links HIV latency with FOXO1 activity via ER stress signaling, but without general T cell activation. Having shown that FOXO1 inhibition prevents latency establishment and also successfully purges the virus from its latent reservoirs, we propose that combining early ART with a FOXO1 inhibitor may downsize the latent reservoir.

## Results

### FOXO1 is a Specific Regulator of HIV Latency Establishment

To test the potential of FOXO1 inhibition to affect HIV-1 latency, we used the second-generation dual-color reporter virus HIV_GKO_ (LTR-HIV-Δ-env-nefATG-csGFP-EF1α-mKO2)^19^ and the FOXO1 inhibitor AS1842856 (Figure 1a). In this system, latently infected K562 cells express the mKO2 fluorescent protein under the control of the EF1α promoter, while cells containing a productive and active virus express both mKO2 and GFP fluorophores, the latter controlled by the HIV-1 LTR. Upon treatment with increasing concentrations of AS1842856, the proportion of latently infected cells decreased, despite no significant changes in the total infection rate or viability (Figure 1a). Concomitantly, the proportion of actively infected cells increased, as expected. The IC_50_ of the inhibitor observed in our cell culture system was 74 ± 15 nM, near its *in vitro* IC_50_ of 33 nM^20^. Thus, FOXO1 inhibition prevented HIV-1 latency establishment and increased the ratio between actively and latently infected cells.

**Figure 1.**
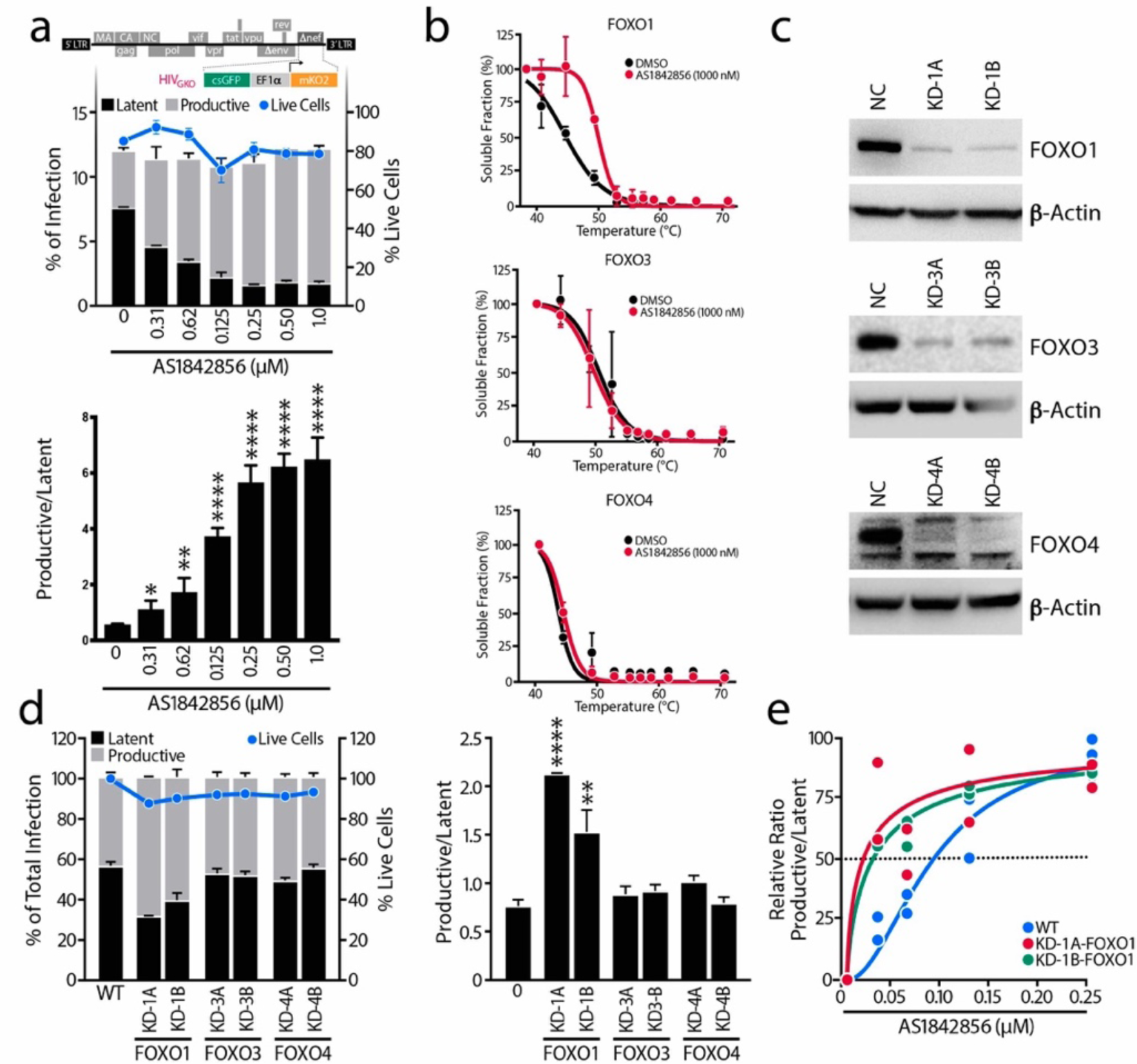
FOXO1 is a Specific Regulator of HIV Latency Establishment. **a**, Schematic representation of HIV_GKO_ dual-labeled HIV-1 reporter. K562 cells were treated with increasing concentrations of the FOXO1 inhibitor AS1842856 just after HIV_GKO_ infection. After 3-4 days, the percentage of latent and productively infected cells were quantified by FACS. In upper panel, amount of productively or latently infected cells (bars) and total cell viability (dots) of a representative experiment. In the lower panel, ratios of productive versus latent populations relative to the total infection rate upon increasing concentrations of AS1842856 treatments. Data are represented as mean ± SD of triplicate values, representative of three independent experiments. **b**, FOXO1, FOXO3 and FOXO4 CETSA-melting curve shifts upon the presence or absence of AS1842856 1,000 nM in K562 cells. Band intensities obtained from western blot analysis were normalized to the highest western blot signal. Relative FOXO-band intensities were plotted against corresponding incubation temperatures and a nonlinear least-squares regression fit was applied. Data represent the mean ± SD of three independent experiments, except for FOXO4 (n=2). **c**, Efficiency of FOXO1, FOXO3 and FOXO4 knockdowns with two different sgRNAs determined by western blot. Cells transduced with NC (negative control) sgRNA lentiviruses were used as a control. **d**, In the left graph, percentage of productive or latent cells relative to the total infection rate and cell viability in the different single knockdown K562 cell lines. In the right graph, ratios of productive versus latent populations in K562 cell lines with knockdown of FOXO1, FOXO3, or FOXO4. Data are represented as mean ± SD of triplicate values, representative of two independent experiments. **e**, Ratios of productive versus latent populations upon increasing concentrations of AS1842856 in the WT K562 or in the FOXO1-knockdown cell lines. Data are represented as mean ± SD of triplicate values, representative of two independent experiments and fitted into a model.

We next sought to confirm that the specificity of AS1842856 for FOXO1. To that end, we performed cellular thermal shift assays (CETSA) against the FOXO isoforms expressed in K562 cells. CETSA is unique in its capability to evaluate biophysical binding under physiological conditions and in living cells. Binding was considered ‘positive’ if there was a significant change in the thermal denaturation profile of the protein upon addition of AS1842856 versus without the drug assessed by western blot (Extended Data Figure 1a). We observed a 5.2 °C shift in FOXO1 melting curve upon AS1842856 treatment (1 μM), whereas no changes were found in either FOXO3 or FOXO4 (Figure 1b), confirming the specificity of the inhibitor. Similar results were observed at 100 nM of AS1842856 (Extended Data Figure 1a-c).

To interrogate the role of other FOXO family members in HIV-1 latency establishment, we generated single knockdowns of FOXO1 (KD-1A-FOXO1, KD-1B-FOXO1), FOXO3 (KD-3A-FOXO3, KD-3B-FOXO3) and FOXO4 (KD-4A-FOXO4, KD-4B-FOXO4) by CRISPR interference (CRISPRi) in K562 cells (Figure 1c). Because FOXO proteins control the cell cycle and promote a quiescent state, each knockdown led to an increase in the proliferation rate (Extended Data Figure 1g), in agreement with previous findings on human bladder cancer cells^21^ and MCF-7 breast adenocarcinoma cells^22^. However, only FOXO1 downregulation increased productive infection and decreased latent infection (Figure 1d). Furthermore, FOXO1 knockdown rendered infected cells more sensitive to AS1842856 treatment, shifting the IC_50_ (Figure 1e and Extended Data Figure 1d-f). Taken together, these data indicate that pharmacological or genetic inhibition of FOXO1, but not FOXO3 or FOXO4, decreases establishment of HIV-1 latency.

### FOXO1 inhibition Reactivates HIV-1 from Latency

The J-Lat clonal cell lines are Jurkat cells that contain a stably integrated, latent HIV-1 (HXB) Δ env provirus with a GFP reporter gene in place of the *nef* gene^23^. In this system, reactivation from established latency can be monitored through GFP expression. After 48 and 72 hours of treatment, AS1842856 stimulated GFP mRNA expression in J-Lat 5A8 cells to a similar degree as TNFα, a known LRA in J-Lat cells, indicating that FOXO1 inhibition can also reverse established latency (Figure 2a). No major effect on cell viability was observed (Figure 2b). Maximal activation by TNFα was seen after 24 hours, while AS1842856-induced activation progressively increased only after this time point.

**Figure 2.**
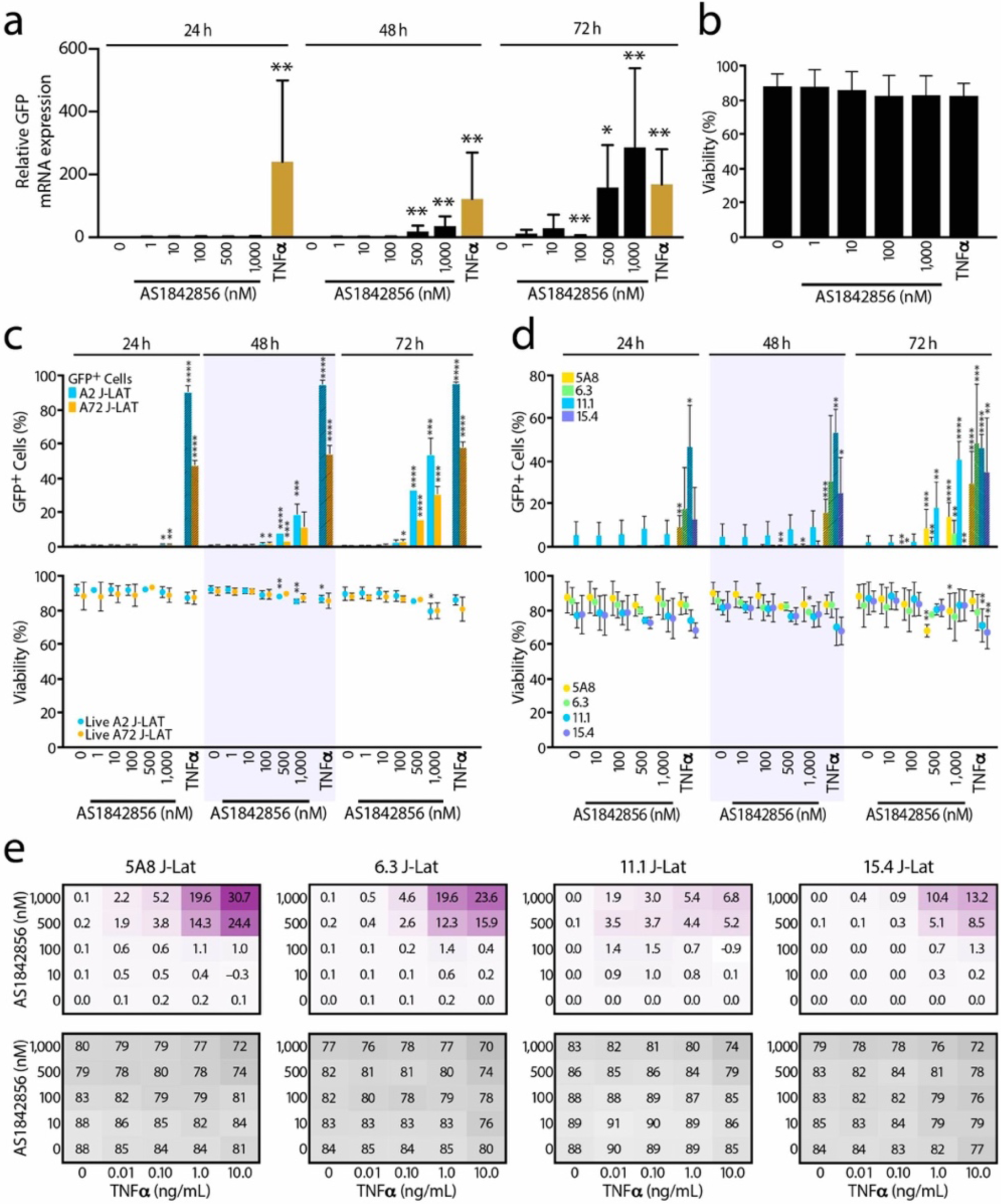
FOXO1 Inhibition Reactivates HIV1 from Latency. **a**, J-Lat cell line 5A8 was treated with increasing concentrations (1-1,000 nM) of AS1842856 for 24, 48 and 72 h and HIV-GFP mRNA reactivation was assessed by RT-qPCR and normalized to *RPL13A* mRNA. Data represent average ± SD of three independent experiments. p-value relative to the control at each time point. **b**, Cell viability assessed by flow cytometry at 72 h of the same experiment as in a. Cell viability was measured by gating on both the live population at the forward scatter (FSC) and side scatter (SSC) plot after staining with the viability dye. Data represent average ± SD of at least three independent experiments. **c**, J-Lat cell lines A2 and A72 were treated with increasing concentrations of AS1842856 for 24, 48 and 72 h and HIV-GFP reactivation (bars) and cell viability (dots) were analyzed by FACS. HIV-GFP reactivation is reported as a percentage of GFP-expressing cells (% GFP+ cells). Data represent average ± SD of at least three independent experiments. **d**, Same experiments as in Fig. 2c but performed in J-Lat cell lines 5A8, 6.3, 11.1 and 15.4. Data represent average ± SD of at least three independent experiments. 10 ng/mL TNFα was used as control. **e**, J-Lat cell lines 5A8, 6.3, 11.1 and 15.4 were treated for 72 h with increasing concentrations of both AS1842856 (Y-axis) and TNFα (X-axis) alone or in combination and analyzed by FACS. Calculation of synergy for drug combinations using the Bliss independence model applied to the HIV-GFP reactivation measured as a percentage of GFP-expressing cells (% GFP+ cells). Synergy (in purple) and cell viability (in grey) data points represent the mean effect from three independent experiments.

Treatment with AS1842856 also increased HIV-driven GFP protein expression in a dose-dependent manner, as determined by flow cytometry (Figures 2c,d). Because FOXO1 and FOXO4 antagonize Tat-mediated transactivation of the HIV-1 promoter through the repression of the viral transactivator Tat^17^, we tested the ability of AS1842856 to reverse HIV latency in J-Lat cells with and without Tat. AS1842856 increased GFP expression in both J-Lat A2 cells, which do contain Tat, and in A72 cells, which do not^23^, indicating that the effect of FOXO1 inhibition on HIV reactivation is Tat-independent (Figure 2c and Extended Data Figure 2a). Again, no effect on cell viability was observed (Figure 2c).

To rule out effects based on clonal variation or proviral integration sites, we compared AS1842856-induced HIV reactivation in four J-Lat clonal cell lines with the latent HIV-1 inserted at distinct genomic sites. After 72 hours of treatment, AS1842856 reactivated HIV in each cell line, as determined by percentage of GFP^+^ cells (Figure 2d) and the median fluorescence intensity (Extended Data Figure 2b). In all cell lines, viability was unaffected, and treatment with TNFα induced GFP expression as early as 24 hours after treatment as expected (Figure 2d). Thus, the effect of AS1842856 is independent from the proviral integration site.

Given the different time courses of TNFα- and AS1842856-induced HIV reactivation, we speculated that these LRAs target different mechanisms of reactivation and might synergize. To test this hypothesis, we co-treated multiple J-Lat lines with increasing concentrations of AS1842856 and TNFα (Extended Data Figure 2c, d). We used the Bliss independence model to assess dose combinations that result in greater than additive effects^24,25^. We observed such synergistic effects across different doses in all clones (Figure 2e, upper panels). Again, we found no major changes in viability in co-treated cell lines (Figure 2e, bottom panels). Collectively, these data show reversal of established latency by FOXO1 inhibition and indicate a molecular mechanism of activation distinct from TNFα.

### FOXO1 inhibition Regulates HIV-1 Latency in Primary CD4^+^ T cells

In order to verify these results in primary T-cell models of HIV latency, we measured the capacity of FOXO1 inhibition to impair HIV latency establishment by infecting resting CD4^+^ T cells from anonymous blood donors with HIV_GKO_ while treating the cells with increasing doses of AS1842856 for 72 hours. As observed previously in K562 cells, inhibition of FOXO1 promoted productive infection and decreased latency (Figure 3a). We also tested the ability of AS1842856 to reverse established HIV latency in primary cells by spin-infecting resting CD4^+^ T cells with a pseudotyped HIV reporter virus containing the firefly luciferase gene^26^. After 6 days in culture, cells reactivated luciferase expression in response to increasing concentrations of FOXO1 inhibitor (Figure 3b) as well as in response to generalized T cell activation induced with either with a combination of phytohemagglutinin and interleukin-2 (PHA/IL-2) or with antibodies directed against the CD3 and CD28 receptors (Extended Data Figure 3b). Notably, FOXO1 inhibition did not induce generalized T cell activation, as measured by expression of CD69 and CD25 activation markers on the cell surface (Extended Data Figure 3a). Combining AS1842856 (100 nM) with a low dose of prostratin (250 nM), a natural protein kinase C activator that induces partial T-cell activation^24,27^, significantly increased the AS1842856 effect in primary T cells, mirroring the synergistic effect of TNFα in J-Lat cells. No change in primary T-cell viability was observed in response to FOXO1 inhibition (Figures 3a-c).

**Figure 3.**
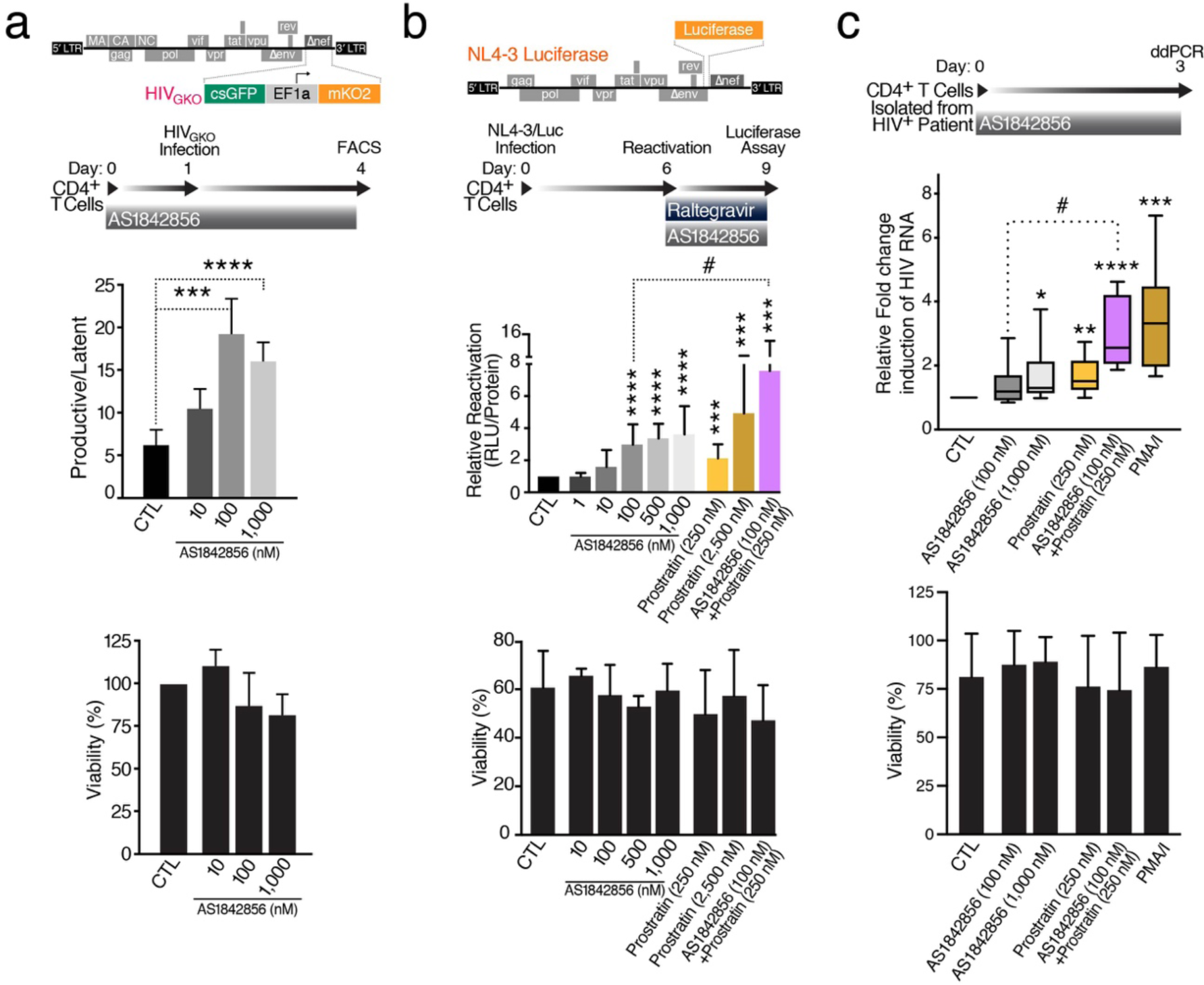
FOXO1 Inhibition Prevents Latency Establishment and Reactivates HIV in Primary CD4^+^ T cells and HIV-infected CD4^+^ T cells. **a**, Schematic representation of HIV_GKO_ dual-labeled HIV-1 reporter and of strategy used to treat primaryCD4^+^ T cells purified from blood of healthy donors and pre-treated for 24 h with increasing concentrations of AS1842856. After infection of HIV_GKO_, resting cells were treated for 3 days with the same amounts of AS1842856 as in the pre-treatment. Top panel shows ratios of productive versus latent populations of infected CD4^+^ T cells from four different healthy donors after treatment. Lower panel shows a histogram plot of percent live cells for each drug treatment relative to the control. Data are represented by mean ± SD of four different donors. **b**, Schematic representation of HIV_NL4-3 Luciferase_ reporter virus and of experimental procedure with primary CD4^+^ T cells. Briefly, CD4^+^ T cells were purified from blood of healthy donors, were infected with HIV_NL4-3 Luciferase_ and, after 6 days, the virus was reactivated for 3 days with increasing concentrations of AS1842856 and prostratin alone or in combination, in the presence of raltegravir (30 μM). HIV reactivation was measured by luciferase activity and cell viability by flow cytometry. Data are mean ± SEM of at least 3 individual donors. **c**, Fold change of cell-associated HIV-1 mRNA expression measured by ddPCR of CD4^+^ T cells of HIV-infected patients on antiretroviral-therapy with undetectable viral load treated with 50 ng/ml (81 nM) PMA + 1 μM Ionomycin (PMA/I) and with 0.2 % DMSO (CTL). AS1842856- and/or prostratin-treated CD4^+^ T cells led to an increase in fold change of cell-associated HIV mRNA expression. Cell viability was assessed by Trypan Blue exclusion. Data are represented as mean ± SD of nine independent experiments. p-value: *p<0.05, **p<0.01, ***p<0.001, ****p<0.0001, relative to CTL (DMSO); #p<0.05, relative to AS1842856 (100 nM).

We replicated these observations in CD4^+^ T cells, isolated from five HIV-infected patients on antiretroviral therapy for at least 6 months and with undetectable viral loads (< 50 copies/ml). Cells were treated *ex vivo* with two concentrations of AS1842856 (100 and 1,000 nM), prostratin (250 nM) or combinations thereof for 3 days, and HIV RNA induction was measured by digital droplet PCR (ddPCR). A two-fold induction was observed with AS1842856 treatment alone, which was significantly enhanced with prostratin cotreatment to similar levels as achieved by PMA/I treatment (Figure 3c). These results confirm a role of FOXO1 in the establishment and maintenance of HIV latency in primary resting T cells.

### Transcriptional Reprogramming in Response to FOXO1 Inhibition

To determine how FOXO1 inhibition changes transcription of resting CD4^+^ T cells, we used RNA sequencing (RNA-Seq) of primary CD4^+^ T cells treated with AS1842856 or DMSO for 48 hours, the first time point at which HIV-1 reactivation was observed. We identified 172 genes significantly up-regulated and 160 genes significantly down-regulated by AS1842856 (q < 0.05) (Figure 4a). Using gene ontology enrichment analysis to integrate the differentially expressed transcripts into biological pathways, we observed significant enrichment for genes involved in the biosynthesis of L-serine (e.g. *PSAT1, PHGDH*), tRNA charging (e.g. *MARS, VARS, GARS*), biosynthesis of purine (e.g. *PFAS, PAICS*) and coenzyme and folate metabolism (e.g. *MTHFD1L, SHMT1*) among the genes upregulated after FOXO1 inhibition. Down-regulated genes included those involved in IL-8 production (e.g. endosomal Toll-like receptors 3, 7 and 8) (Figure 4b).

**Figure 4.**
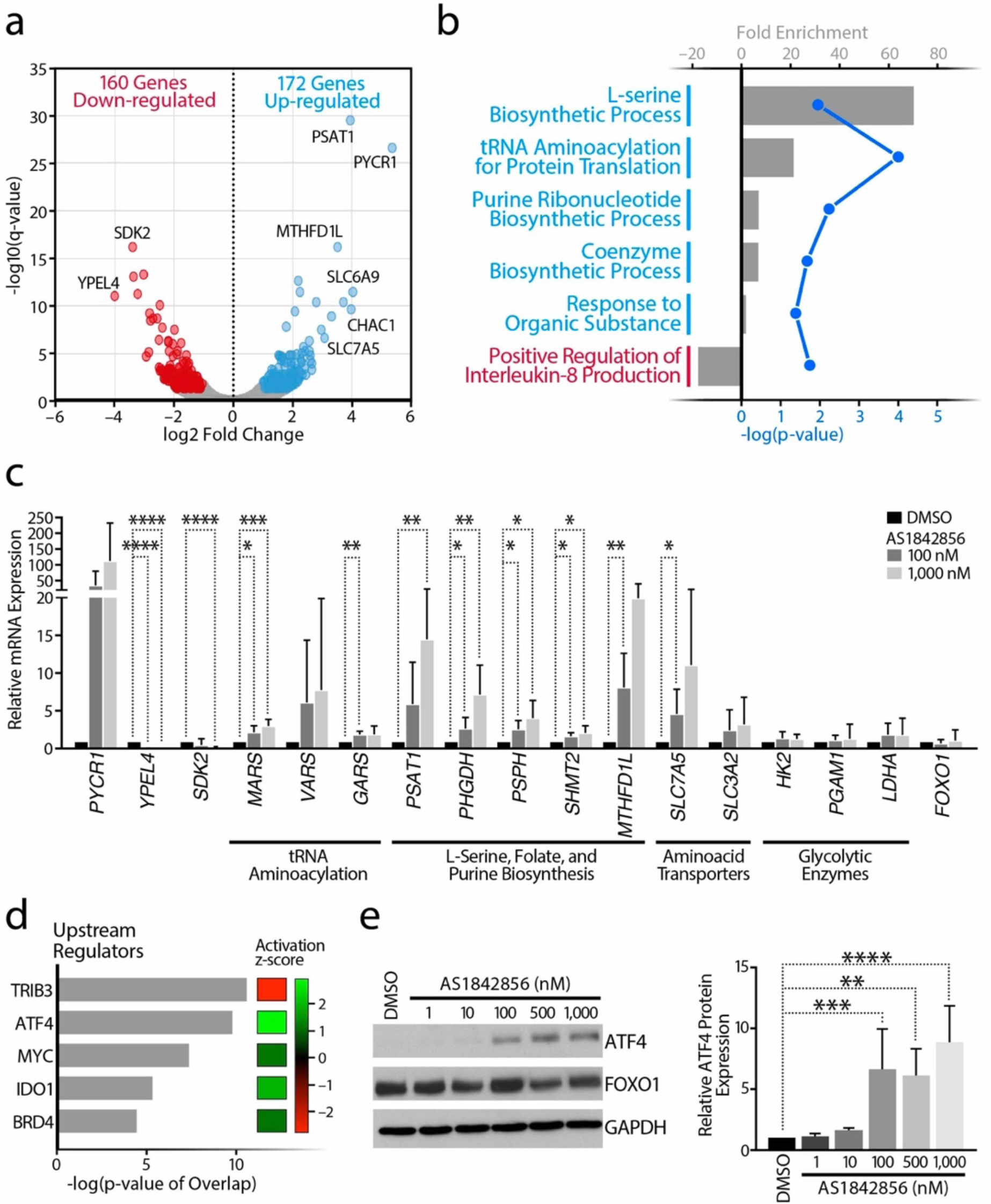
Marked Upregulation of the Regulator ATF4 in Response to FOXO1 Inhibition in Primary CD4^+^ T cells. **a**, Volcano plot from the RNA-Seq data comparing CD4^+^ T cells treated with AS1842856 (1,000 nM) versus DMSO control for 48 h. Up-regulated (blue) or down-regulated (red) genes are q-value < 0.05 and log_2_ fold change ≥ 1 or ≤ −1, respectively. **b**, GO pathway analysis of the most dysregulated canonical pathways for the up- and down-regulated genes. **c**, Confirmation of up- or down-regulated expression after 48 h treatment of specific genes from altered pathways by RT-qPCR, normalized to *RPL13A* mRNA. Data represent mean ± SD of six individual donors. **d**, Analysis of the five top upstream regulators and table with their activation z-scores according to the RNA-Seq data. **e**, Representative blot of protein expression of the transcription factors ATF4 and FOXO1 (left) and densitometry analysis of ATF4 protein expression (right) was performed from three individual donors. Data are mean ± SD.

The RNA-Seq results were validated by measuring read counts of the selected genes representative of different pathways (Extended Data Figure 4d) as well as by real-time quantitative PCR (Figure 4c). Although FOXO1 has been connected with the prevention of glycolysis^28^, treatment with AS1842856 did not affect expression of glycolytic enzymes or FOXO1 itself. Upstream regulator analysis identified TRIB3 and ATF4 as the top regulators of AS1842856-induced gene changes (Figure 4d). Specifically, the algorithm predicted that TRIB3 was highly inhibited, while ATF4 was activated (Figure 4d). TRIB3 is known to inhibit ATF4^29,30^, a transcription factor specifically upregulated at the protein level as part of the integrated stress response, which involves phosphorylation of the translation initiation factor eIF2α by various stress-specific kinases^31–33^. RNA-Seq analysis of AS1842856-treated CD4^+^ T cells at 12 hours after treatment showed upregulation of pathways including ribosomal large subunit assembly, rRNA processing, and the unfolded protein response (UPR), supporting a role for ATF4 and the integrated stress response in gene changes after FOXO1 inhibition (Extended Data Figure 4a-c). In line with this hypothesis, FOXO1 inhibition increased expression of ATF4 protein levels in CD4^+^ T cells across multiple donors (Figure 4e).

### FOXO1 Inhibition Induces HIV-1 Reactivation via ATF4 and NFAT

Because ATF4 binds the SIV and HIV LTR^34^ and activates the related human T-cell leukemia virus type 1 (HTLV-1)^35^, we performed chromatin immunoprecipitation (ChIP) experiments to test whether ATF4 binds the HIV LTR in response to FOXO1 inhibition. While low levels of ATF4 bound HIV LTR chromatin in untreated J-Lat cells, its recruitment was 3-fold enriched after AS1842856 treatment (Figure 5a). We also observed a marked increase in RNA polymerase II recruitment to the HIV promoter after FOXO1 inhibition, consistent with HIV transcription reactivation. The RelA subunit of the NF-κB transcription factor, a key regulator of HIV transcription in response to T cell activation or TNFα exposure^36^, was not recruited to the HIV LTR after AS1842856 treatment but was significantly enriched after TNFα treatment, as expected (Figure 5a). Nuclear factor of activated T cells (NFAT) can occupy the NF-κB binding sites in the HIV LTR^37^, and is activated through calcium mobilization after T-cell activation, rather than by protein kinase C activation, as is the case for NF-κB^38,39^. NFAT was markedly enriched at the HIV LTR after FOXO1 inhibition (Figure 5a), indicating that FOXO1 in resting T cells suppresses a (non-canonical) transcriptional program that controls HIV transcription without NF-κB, but with ATF4 and NFAT. This result explains the synergy observed between AS1842856 and TNFα or prostratin, as these compounds activate HIV through NF-κB^27^.

**Figure 5.**
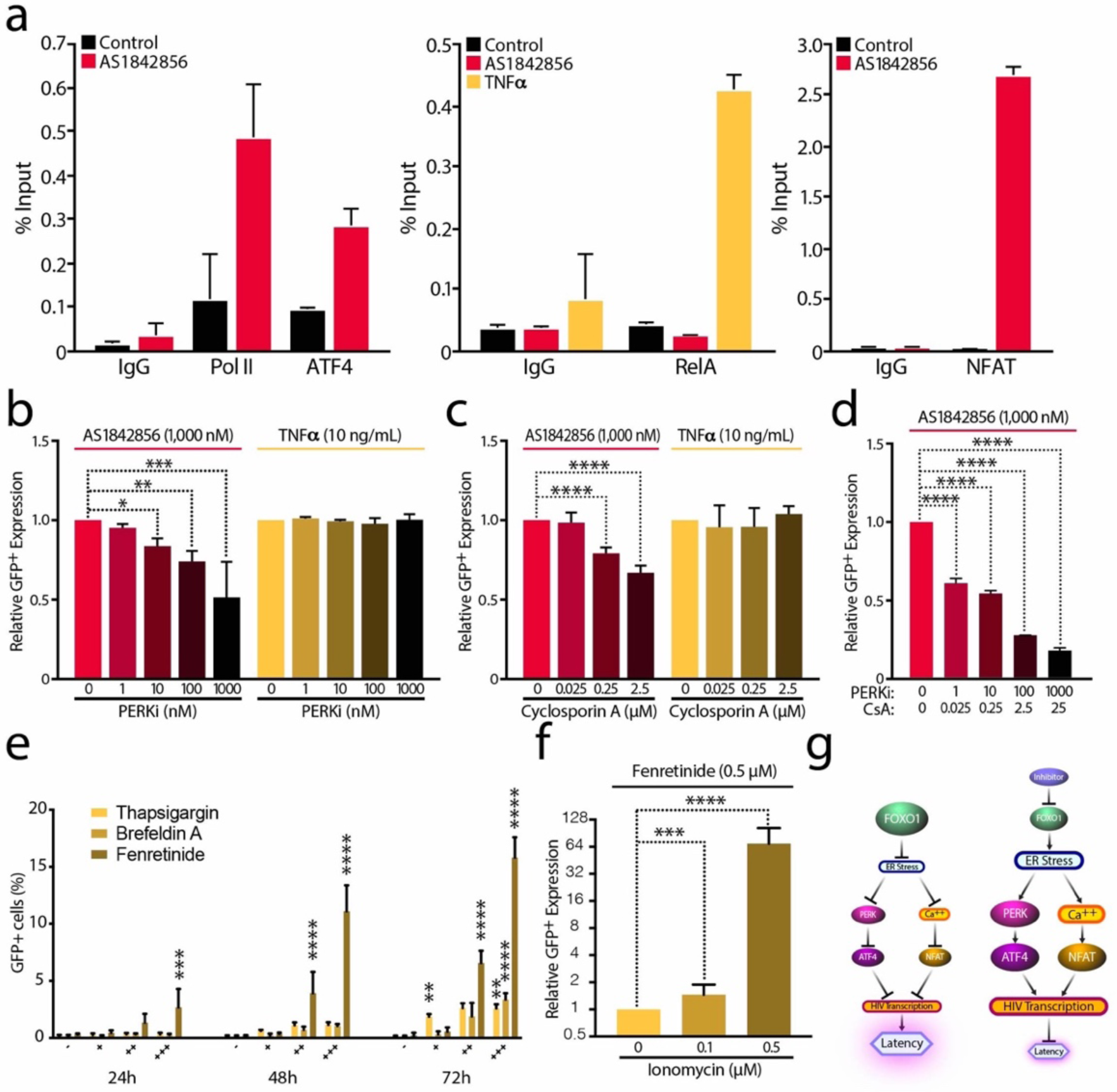
FOXO1 Inhibition Induces HIV Reactivation in the Absence of NF-kB Recruitment via ATF4 and NFAT. **a**, Chromatin immunoprecipitation (ChIP) assays with antibodies against Pol II, ATF4, RelA, NFAT and IgG control at the HIV LTR, followed by qPCR using primers specific for HIV-1 LTR Nuc0 or Nuc1. Chromatin was prepared from J-Lat A2 and 5A8 cells, in which the LTR was stimulated by 1,000 nM AS1842856 treatment, 10 ng/mL TNFα or which were left untreated/DMSO. Representative experiment of at least three independent experiments. **b**, J-Lat cell line A58 pre-treated for 1 h with increasing concentrations of PERKi (GSK2656157 / PERK inhibitor II) before being treated with 1,000 nM AS1842856 (72 h) or 10 ng/mL TNFα (24 h). HIV-GFP reactivation was analyzed by FACS and normalized to the control. Data shown are mean ± SD of three independent experiments. **c**, Same experiment as in Fig. 5b, but cells were treated with increasing concentrations of Cyclosporin A (CsA), and without 1 h pre-treatment. **d**, Similar experiment that in Fig. 5b or Fig. 5c, but combining increasing concentrations of PERKi (GSK2656157 / PERK inhibitor II) and Cyclosporin A (CsA). Data represent average ± SD of at least three independent experiments. **e**, J-Lat cell line A58 was treated with increasing concentrations of Thapsigargin (0.01, 0.1, 1 µM), Brefeldin A (0.01, 0.1, 1 µg/mL) and Fenretinide (0.5, 2, 5 µM) for 24, 48 and 72 h and HIV-GFP reactivation was analyzed by FACS. **f**, J-Lat cell line A58 was treated with 0.5 µM Fenretinide and increasing concentrations of Ionomycin. HIV-GFP reactivation and cell viability were analyzed by FACS. Data are mean ± SD of three independent experiments. **g**, Model: FOXO1 inhibition leads to ER Stress. Thus, ATF4 activation through PERK and NFAT via cytosolic calcium release will promote HIV transcription and will prevent HIV latency.

To investigate integrated stress response’s potential role in HIV reactivation, we evaluated the effect of AS1842856 in combination with inhibitors of different eIF2α upstream kinases: the double-stranded RNA-dependent protein kinase R (PKR), activated by viral infection; the PKR-like endoplasmic reticulum kinase (PERK), activated in response to ER stress; and the general control nonderepressible 2 (GCN2) kinase, activated by amino acid deficiencies^40–45^. Treatment with a pharmacological inhibitor of PERK, but not GCN2 or PKR, reduced AS1842856-induced HIV reactivation, consistent with the model that FOXO1 inhibition activates ATF4 through ER stress and the integrated stress response (Figure 5b and Extended Data Figure 5a,b). In contrast, TNFα-mediated HIV reactivation was not affected by PERK inhibition (Figure 5b) but was diminished upon GCN2 or PKR inhibitor treatment (Extended Data Figure 5a,b). This finding underscores the different molecular mechanisms underlying latency reversal by the two activators.

Because NFAT was also recruited to the HIV LTR after FOXO1 inhibition but is not regulated through PERK activation, we tested the effect of cyclosporin A (CsA), a potent inhibitor of calcium release and NFAT mobilization known to reduce HIV transcriptional activity^46^. Increasing concentrations of CsA suppressed HIV reactivation induced by AS1842856, but not TNFα, supporting a specific role of NFAT in FOXO1 inhibitor-mediated latency reversal (Figure 5c). Treatment with the PERK inhibitor or CsA alone suppressed HIV reactivation by ∼ 42.5 % (49 ± 22 % (PERKi); 36 ± 7 % (CsA)), whereas the combination almost completely blocked AS1842856-mediated viral reactivation (83 ± 4 %) (Figure 5d). This result suggests that both ATF4 and NFAT are important for HIV reactivation in response to FOXO1 inhibition.

Both transcription factors are linked to ER stress, either through activation of PERK or mobilization of ER-resident calcium stores^47,48^. To test the concept that ER stress reactivates HIV from latency, we treated J-Lat cells with known inducers of ER stress^49^, including thapsigargin, a sarco/ER Ca^2+^ ATPase [SERCA] inhibitor, Brefeldin A, a retrograde vesicular transport inhibitor, and fenretinide, a clinically tested synthetic retinoid derivative. All resulted in HIV-1 reactivation, and fenretinide treatment acted most strongly (Figure 5e). Activation by these agents was associated with dose-dependent cellular toxicity, explained by their known function as potent apoptosis inducers^50^ (Extended Data Figure 5c). To minimize toxicity caused by high concentrations of fenretinide, we combined a nontoxic dose of fenretinide (0.5 μM) with increasing amounts of ionomycin to mobilize additional Ca^2+^ ions. Although ionomycin alone was sufficient to induce HIV reactivation (Extended Data Figure 5d), the combination significantly increased the effect (∼64-fold over fenretinide alone) with minimal cellular toxicity (Figure 5f and Extended Data Figure 5e). These data support a model where FOXO1 interferes with potentially low-level continuous ER stress in resting T cells, suppressing activation of ATF4 and NFAT, thereby promoting HIV latency. Inhibition of FOXO1 activity unleashes ER stress signaling and calcium release, mobilizing the two transcription factors, which then activate the HIV-1 LTR and reverse viral latency (Figure 5g).

## Discussion

By uncovering a link between FOXO1 and ER stress, this work describes a new mechanism through which HIV latency is established and maintained in quiescent CD4^+^ T cells. Our results are in agreement with work linking ER stress with CD4^+^ T cell activation^40,51^, and showing that FOXO activity protects quiescent cells from oxidative stress^52^. We extend these studies to show that this effect does not involve NF-κB activity, but that FOXO1 activity controls ATF4 and NFAT mobilization. In other systems, ATF4 interacts with FOXO1 and regulates glucose metabolism in osteoblasts^53^, while FOXO activity suppresses calcineurin/NFAT activation by reducing oxidative stress in cardiac physiology^54,55^.

Malfunctioning of the ER, like excessive secretory activity, Ca^2+^ depletion, or presence of misfolded proteins, has a pivotal role in the life cycle of viruses establishing latent infections. ER stress activates the unfolded protein response which is necessary for replication of many viruses, such as herpes viruses^56^. Other viruses are known to hijack the unfolded protein response to promote the expression of ATF4, like hepatitis C virus^57^, West Nile virus^58^, respiratory syncytial virus^59^, Japanese encephalitis virus^60^, human cytomegalovirus^61^, dengue virus (DENV)^62^, or infectious bronchitis virus (IBV)^63^. In the context of HIV, our work shows that ER stress promotes active HIV infection with FOXO1 acting as a critical regulator of this process in resting T cells. These results agree with previous findings that HIV-1 infection is associated with enhanced expression of the ER stress sensors PERK, ATF6, and IRE-1, in PBMCs^64^, and that the viral protein trans-activator Tat induces ER stress in astrocytes^65^.

While NFAT is not a “classical” ER stress-induced factor, it was one of four new transcription factors found activated by ER stress^47^. In addition, its close connection with intracellular calcium levels links it to the ER, where the majority of intracellular calcium is stored^66^. Prolonged ER stress is associated with calcium release, and the ER-resident chaperone binding immunoglobulin protein (BiP), an important sensor of unfolded proteins, is itself calcium-regulated^67^. It remains unknown what causes ER stress, unleashed after FOXO1 inhibition, in quiescent T cells, but every protein that enters the ER can potentially trigger a response. PERK is one of three sensors in the ER and when nonactivated it is maintained in a monomeric state, bound to BIP. When BiP senses the accumulation of mis-folded proteins, it dissociates from PERK, allowing the kinase to dimerize and trans-autophosphorylate^68^. Phosphorylated PERK promotes phosphorylation of eIF2α, which impairs translation of most mRNAs, with the exception of a few transcripts like ATF4 that are specifically expressed because of the presence of multiple open reading frames in the 5’ untranslated regions^69^.

Further experiments are required to clarify the potential involvement of ATF4-induced downstream pathways in HIV reactivation and to better understand the molecular link between FOXO1, ER stress and proteostasis. PERK phosphorylates FOXO1 to enhance insulin responsiveness^70^ but no effect of FOXO1 on PERK activity has yet been reported. Nevertheless, the discovery of this new mechanism opens the door to promising therapeutic options, especially considering the synergistic effects of FOXO1 inhibition in combination with other LRAs and the latency reversing effect of ER stress-inducing agents such as fenretinide, currently in early clinical trials for cancer^71^. An important aspect is that the cytolytic activity of CD8^+^ T cells, key contributors to a successful “shock and kill” strategy, is strengthened by treatment with the FOXO1 inhibitor^15^, in contrast to HDAC inhibitors, which suppress CD8^+^ T cell activity^72^. A combination of FOXO1 inhibitors with early ART seems especially promising, given the negative effect on latency establishment, potentially prolonging the time window for intervention to prevent maximal reservoir establishment.

## Methods

### Cell lines and primary cultures

J-Lat cell lines (clones A2, A72, 6.3, 11.1 15.4 and 5A8) were obtained from the Verdin and Greene Laboratories at the Gladstone Institutes, which generated these lines from Jurkat cells^23,73^. J-Lat cells were cultured at 37°C in RPMI (Corning, VA, USA), supplemented with 10% FBS (Corning, VA, USA), 1% L-glutamine (Corning, VA, USA) and 1% penicillin-streptomycin (Corning, VA, USA). K562 cell lines expressing dCas9-BFP-KRAB were a gift from the lab of Johnathan Weissman, at UCSF^74^ and were maintained in appropriate volume of RPMI medium supplemented with 10% fetal bovine serum (Serum Plus - II, Sigma MO, USA) and 1% penicillin-streptomycin. HEK293T cells (ATCC) were cultured in DMEM (Corning, VA, USA), supplemented with 10% FBS, 1% glutamine, and 100 U/mL penicillin-streptomycin at 37°C, 5% CO_2_. Short-term passages (<15) were used for all experiments. Primary CD4^+^ T cells were isolated from peripheral blood mononuclear cells (PBMCs) and enriched via negative selection with an EasySep Human CD4^+^ T Cell Isolation Kit (Stemcell Technologies, Canada). Primary CD4^+^ T cells were cultured at 37°C in RPMI supplemented with 10% FBS, 1% L-glutamine and 1% penicillin-streptomycin.

### Chemicals

Cells were treated with the following compounds: FOXO inhibitor / AS1842856 (344355, Calbiochem); TNFα (300-01A, PeproTech); Raltegravir (CDS023737, Sigma-Aldrich); Prostratin (P4462, LC Laboratories); PHA-M (10576015, Sigma-Aldrich); IL-2 (I2644, Sigma-Aldrich); αCD3 (40-0038, Tonbo Biosciences); αCD28 (70-0289, Tonbo Biosciences); PMA (P8139, Sigma-Aldrich); Ionomycin (I0634, Sigma-Aldrich); PERK inhibitor II / GSK2656157 (504651, Sigma-Aldrich); GCN2 inhibitor / A-92 (2720, Axon Medchem); Imidazolo-oxindole PKR inhibitor C16 (I9785, Sigma-Aldrich); Cyclosporin A (C3662, Sigma-Aldrich); Thapsigargin (T9033, Sigma-Aldrich); Brefeldin A 1,000X Solution (00-4506-51, ThermoFisher Scientific); Fenretinide (17688, Cayman Chemicals).

### Virus production

Pseuodtyped HIV_GKO_ and HIV_NL4-3/Luciferase_ viral stocks were generated by co-transfecting (standard calcium phosphate method) HEK293T cells with a plasmid encoding HIV_GKO_ or HIV_NL4-3/Luciferase_, and a plasmid encoding HIV-1 dual-tropic envelope (pSVIII-92HT593.1) or vesicular stomatitis virus G protein (VSVg), respectively. Medium was changed 6–8 h after transfection, and supernatants were collected after 48-72 hours, centrifuged (20 min, 2000 rpm, RT), filtered through a 0.45 μm membrane to clear cell debris, and then concentrated by ultracentrifugation (22,000 g, 2 h, 4 °C). Virus-like particles (VLPs) were similarly generated as described previously^75^. Here, HEK293T cells were co-transfected with 16 μg of a plasmid encoding Vpr-Vpx (PSIV3^+^), 16 μg of a plasmid encoding gp160, 40 μg of the packaging vector plasmid 8.91 and 4 μg of pAdvantage DNA (T175 flask). Concentrated virions were resuspended in complete media and stored at −80 °C. Virus concentration was estimated by p24 titration using the FLAQ assay^76^ (HIV_GKO_ and VLPs) or the Lenti-X™ p24 Rapid Titer Kit (Clontech; HIV_NL4-3/Luciferase_).

### Cell infection

K562 cell lines expressing dCas9-BFP-KRAB were spinoculated for 2 h at 2000 rpm at 32°C with the HIV_GKO_ VSV-G pseudotyped dual-reporter virus encoding for GFP and mKO2 (MOI = 0.01), followed by *in vivo* culture with complete RPMI medium. Purified CD4^+^ T cells isolated from healthy peripheral blood cells were also spinoculated with HIV_NL4-3_ VSV-G pseudotyped virus encoding a luciferase reporter at a concentration of 0.25-1 μg of p24 per 1×10^6^ cells. For Figure 3a, purified resting CD4^+^ T cells were infected with VLPs at day 0. Cells were spin-infected in a 96 well V-bottom plates (1×10^6^ cells/well) with 100 ng of p24 VLPs per 1×10^6^ cells (final volume 100 μl/well) for 2 h at 2,000 rpm at 4 °C and then incubated at 37°C. The following day, 100 μL of fresh complete 1640 RPMI with or without the different drugs was added to each well. Finally, 24h later, treated and untreated cells were spin-infected with HIV_GKO_ as described previously^77^. Briefly, 1×10^6^ cell were spin-infected with serial dilutions of HIV_GKO_ (1/2 dilutions), starting with 300 ng of p24 for 2 h at 2,000 rpm at 32 °C. After spin-infection, fresh drugs were added to the media, and cells were cultured for an additional 4 days. Infected cells were analyzed by flow cytometry 4 days post-infection. Resting CD4^+^ T cells were treated 24h pre-HIV**GKO** infection with 0-10-100-1,000 nM AS1842856. The same drug concentrations were added onto cells right after HIV**GKO** spin-infection and were not washed away until data analysis.

### T-cell activation analysis and Flow cytometry

Cells were seeded in drug-supplemented media in triplicates onto 96-well plates with a target cell density of 0.125-0.25-0.5 × 10^6^ cells/mL, and then incubated at 37 °C for 72-48-24 h, respectively. In preparation for flow cytometry, cells were washed in V-shaped 96-well plates (500 x g, 5 min, 4 °C), and stained with Fixable Viability Dye eFluor 780 (65-0865-14; ThermoFisher Scientific). For T cell activation analysis, CD69 and CD25 expression was measured by flow cytometry gating on CD3^+^CD4^+^ T cells using FITC-labeled antibodies for CD3 (11-0048-42, eBioscience), APC-conjugated CD25 antibodies (17-0259-42, eBioscience), PerCP-labeled antibodies for CD4 (300528, Biolegend), and CD69-V450 (560740, BD Horizon). Staining was performed for 30 min on ice in FACS buffer (PBS, 2% FBS) before analysis. 5A8 J-Lat cells were measured on a BD FACSCalibur system (BD Biosciences) in an uncompensated setting. Infected or uninfected K562 and CD4+ T cell data was collected using the BD LSRII flow cytometer (BD Biosciences). Data were analyzed using FlowJo V10.1 software (Tree Star).

### Viability

Cell viability (% survival) was measured by flow cytometry gating on forward and side scatter analysis and using Fixable Viability Dye eFluor 780 (65-0865-14; ThermoFisher Scientific). Trypan blue dye exclusion was used to evaluate the cell viability in the CETSA assays in K562 cells and in the reactivation studies in HIV+ donors. A 10 μL cell aliquot from each sample was briefly mixed with an equal volume of 0.4% (w/v) trypan blue dye solution (Gibco, NY, USA) and counted. Cells with the ability to exclude trypan blue were considered viable.

### Cellular Thermal Shift Assay (CETSA)

For CETSA, K562 cell were freshly seeded one day before the experiment. On the day of the experiment, equal numbers of cells were counted by Moxi Mini automated cell counter (Orflo) and 0.6×10^6 cells per data point were seeded in T-25 cell culture flasks (VWR, PA, USA) in appropriate volume of culture medium. Cells were exposed to 100 nM AS1842856 or equal DMSO volume for 3 hours in an incubator with 5% CO2 and 37°C. Following the incubation, cells were harvested, washed with PBS and diluted in PBS supplemented with EDTA-free complete protease inhibitor cocktail (ROCHE). Then, the cells were divided into 100 μl aliquots and heated individually at different temperatures for 3 minutes (Thermal cycler, BIO-RAD) followed by cooling for 2 minutes at room temperature. Cell suspensions were freeze-thawed three times with liquid nitrogen. The soluble fraction was separated from the cell debris by centrifugation at 20000 g for 20 minutes at 4°C. The supernatants were transferred to new microcentrifuge tubes and analyzed by SDS-PAGE followed by immunoblotting analysis. CETSA experiments were performed as triplicates on different days. Immunoblotting results were subjected to densitometric analysis (Image J) and melting temperatures of FOXO1, FOXO3 and FOXO4 were determined by Prism 7 (GraphPad) using nonlinear least-squares regression fit to Y = 100/ (1 + 10 ^ (Log IC_50_ – X) *H), where H = Hill slope (variable). The viability of the cells was assessed in triplicate by trypan blue exclusion.

### Generation of stable CRISPRi Knockdown

Individual sgRNAs from human v2 library^78^ were cloned into lentiviral vectors expressing either a Puromycin resistance cassette (60955, Addgene) or into a modified vector expressing a Blasticidin resistance cassette (Gift from Nevan Krogan, UCSF). Lentiviruses were prepared in HEK293T. For single knockdowns, K562 cell lines expressing dCas9-BFP-KRAB were transduced with a single construct and selected with either Puromycin (1 μg/mL, Sigma-Aldrich) or Blasticidin (10 μg/mL, Sigma-Aldrich). See Table 1.

### Western blot

Cells were lysed in radioimmunoprecipitation assay buffer (50 mM Tris-HCl, pH 8, 150 mM NaCl, 1 % NP-40, 0.5 % sodium deoxycholate, and 0.1 % SDS, supplemented with protease inhibitor cocktail; Sigma-Aldrich) for 30 min at 4°C and the protein content was measured by DC Protein Assay (Bio-Rad). Protein samples (20 μg) were resuspended in Laemmli buffer and separated by SDS-PAGE. Next, samples were transferred to PVDF membranes (Immobilon-P, Millipore) and blocked with 5 % dry milk supplemented with 0.05% Tween 20 in PBS. The membranes were then immunoblotted by specific antibodies: FOXO1 (2880; Cell Signaling Technology), FOXO3 (D19A7; Cell Signaling Technology), FOXO4 (9472; Cell Signaling Technology), ATF4 (11815; Cell Signaling), β-actin (A5316; Sigma-Aldrich) and GAPDH (5174; Cell Siganling). For chemiluminescent detection, we used enhanced luminol-based chemiluminescent substrate (ECL) and ECL Hyperfilm (Amersham). Immunoblotting results were analyzed by densitometry (Image J).

### RNA extraction, RT, and quantitative RT-PCR

Total RNA from samples was extracted using the Direct-zol RNA kit (R2060; Zymogen). cDNA was generated using 500 ng for CD4^+^ T cells or 1,000 ng of total RNA for J-Lats with Superscript III reverse transcription (18080-044; ThermoFisher) and oligo(dT)_12-18_ (18418-012; ThermoFisher). Quantitative RT-PCR was done using Maxima SYBR Green qPCR Master Mix (Thermo Scientific) on SDS 2.4 software (Applied Biosystems) in a total volume of 12 μL. The SYBR green qPCR reactions contained 5 µl of 2× Maxima SYBR green/Rox qPCR Master Mix (K0221; ThermoFisher), 5 µl of diluted cDNA, and 1 nmol of both forward and reverse primers. The reactions were run as follows: 50°C for 2 min and 95°C for 10 min, followed by 40 cycles of 95°C for 5 s and 62°C for 30 s. See Table S2 for a qRT-PCR primer list. Primer efficiencies were around 100%. Dissociation curve analysis after the end of the PCR confirmed the presence of a single and specific product. All qPCRs were independently repeated at least three times, averaged and compared using standard deviation (±SD).

### Measurement of intracellular HIV-1 mRNA by digital droplet PCR from isolated CD4+ T cells from ART-suppressed individuals

This study sampled HIV-infected participants from the Zuckerberg San Francisco General Hospital clinic SCOPE cohorts. These individuals met the criteria of having had started ART during chronic infection and being under suppressive ART, with undetectable plasma HIV-1 RNA levels (<50 copies/mL) for a minimum of one year. The UCSF Committee on Human Research approved this study, and the participants gave informed, written consent before enrollment. Peripheral blood mononuclear cells (PBMCs) were isolated from fresh blood (100 mL) using Lymphocyte Separation Medium (25-072-CI, Corning) and CD4+ T cells were isolated using negative selection by EasySep kit (19052, STEMCELL) according to the manufacturer’s instructions. Isolated CD4+ T cells were aliquoted at a density of 1×10^6 cells per well in 5 mL RPMI medium supplemented with 10% FBS and corresponding drugs. All drugs were prepared in the culture medium from stock solutions dissolved in DMSO. After the 72 h treatment, total RNAs from the cells were extracted miRNeasy kit (217004, QIAGEN) with the optional on-column DNase treatment step, followed by a TURBO™DNase (AM2238, Invitrogen) treatment post isolation. RNA was quantified using a NanoDrop Spectrophotometer ND-1,000 (NanoDrop Technologies). The SuperScript III RT First-Strand Synthesis system (18080051, Invitrogen) was used to reverse-transcribe 450-1,000 ng of RNA with random hexamers (48190–011, Invitrogen). Following cDNA synthesis, absolute quantification of the HIV Long Terminal Repeat (LTR) was performed in a duplex-digital droplet PCR reaction using the Raindrop System (Raindrop Technologies / Bio-Rad). The Raindrop Source instrument was used to generate uniform aqueous droplets (5 picoliter) for each sample on a 8 well microfluidic chip (Raindrop Source Chip). Cell-associated HIV-1 RNA was detected using LTR-specific primers F522-43 (5’ GCC TCA ATA AAG CTT GCC TTG A 3’; HXB2 522–543) and R626-43 (5’ GGG CGC CAC TGC TAG AGA 3’; 626–643). Reactions were carried out in 50 μl volumes with 2μl 25x Droplet Stabilizer (Raindance Technologies), 25μl 2x Taqman Genotyping Master Mix (Thermo Fisher Scientific), 900nm primers, 250nm probes and 5ul cDNA. Droplets were thermocycled at 95°C for 10 minutes, 45 cycles of 95°C for 15 second, 59°C for 1 minutes, followed by 98°C for 10 minutes. The thermal-cycled 8-tube strip was placed into the deck of the RainDrop Sense instrument with a second microfluidic chip (RainDrop Sense chip) used for single droplet fluorescence measurements. Data were analyzed using the Raindrop Analyst Software on a two dimensional histogram with FAM intensity on the X-axis and Vic intensity on the Y-axis, normalized to RNA input. Each drug combination was normalized to the DMSO (0.2%) control to calculate the fold induction.

### Luciferase Reporter Assay

For reactivation of latent HIV-1 provirus, cells were counted and collected as pellets by centrifugation at 1500 rpm for 10 min. Cells were then plated in 96-well U-bottom plates at 1×10^6^ per 200 μl in the presence of 30 μM Raltegravir (Santa Cruz Biotechnology) and the indicated activator. Cells were harvested 72 h after stimulation, washed one time with PBS, and lysed in 60 μl of Passive Lysis Buffer (Promega). After 15 min of lysis, the luciferase activity in cell extracts was quantified with a SpectraMax i3x Multi-Mode plate reader (Molecular Devices, USA) after mixing 20 μl of lysate with 100 μl of substrate (Luciferase Assay System-Promega). Relative light units (RLU) were normalized to protein content determined by DC Protein assay (BIO-RAD).

### Synergy Bliss model

To analyze drug combinatorial effects, we used the Bliss independence model from the SynergyFinder web application (https://synergyfinder.fimm.fi)^79^. The Bliss score (Δf_axy_) is the difference between the calculated reactivation value if the two drugs act independently and the observed combined reactivation values. Synergy is defined as Δf_axy_ > 0, while Δf_axy_ < 0 indicates antagonism. Positive Bliss scores represent dose combinations which have greater than additive effect^25^.

### RNA-Seq and Ingenuity Pathway Analysis

RNA was prepared from CD4^+^ T cells using the QIAgen RNeasy Plus Kit. The Gladstone Institutes Genomics Core carried out the downstream processing of the RNA samples. Strand-specific cDNA libraries were prepared using the Nugen Ovation kit (Nugen) and the libraries were deep sequenced on NextSeq 500 using single-end 75 bp sequencing. RNA-seq analysis was done using the Illumina RNAexpress application v 1.1.0. Briefly, alignment of RNA-Seq reads was performed with the STAR aligner and, after alignment of aligned reads to genes, differential gene expression was analyzed with DESeq2. Pathway analysis was carried out using Ingenuity Pathway Analysis and the Gene Ontology AmiGO Term Enrichment tool. For the Ingenuity analysis, the entire dataset was uploaded, and a two-direction analysis carried using filters to restrict the analysis to differentially expressed genes as above. Refer to the manufacturer’s website (http://www.ingenuity.com) for further details. For Gene Ontology, individual lists of genes were uploaded and analyzed separately.

### Chromatin immunoprecipitation (ChIP)

J-Lat A2, A72 and 5A8 cells were treated with TNFα (10 ng/ml) or AS1852856 (1,000 nM) for 72 h. Cells were fixed with 1% formaldehyde (v/v) in fixation buffer (1 mM EDTA, 0.5 mM EGTA, 50 mM HEPES, pH 8.0, 100 mM NaCl), and fixation was stopped after 30 min by addition of glycine to 125 mM. The cell membrane was lysed for 15 min on ice (5 mM Pipes, pH 8.0, 85 mM KCl, 0.5% NP40, protease inhibitors). After washing with nuclear swell buffer (25 mM HEPES, pH 7.5, 4 mM KCl, 1 Mm DTT, 0.5% NP-40, 0.5 mM PMSF) and micrococcal nuclease (MNase) digestion buffer (20 mM Tris pH 7.5, 2.5 mM CaCl_2_, 5 mM NaCl, 1 mM DTT, 0.5 % NP-40), the pellet was resuspended in MNase buffer (15 mM Tris-HCl, pH 7.5, 5 mM MgCl_2_, 1 mM CaCl_2_, 25 mM NaCl). Subsequently, samples were incubated with MNase (New England Biolabs) for 30 min at 37°C. The reaction was quenched with 0.5 M EDTA and incubated on ice for 5 min. Nuclei were lysed (1% SDS, 10 mM EDTA, 50 mM Tris-HCl, pH 8.1, protease inhibitors), and chromatin DNA was sheared to 200–1,000 bp average size through sonication (Ultrasonic Processor CP-130, Cole Parmer). Cellular debris was pelleted, and the supernatant was recovered. Lysates were incubated overnight at 4°C with 5 μg antibody against RNA Pol II (39097, Active Motif), RelA (A301-824A, Bethyl), ATF4 (11815, Cell Signaling), NFAT1 (ab2722, Abcam), or IgG control (P120-101, Bethyl). After incubation with protein A/G Dynabeads for 2 h and washing three times with low salt buffer (0.1% SDS, 1% Triton X-100, 2 mM EDTA, 20 mM Tris-HCl, pH 8.1, 150 mM NaCl), one time with high salt buffer (0.1% SDS, 1% Triton X-100, 2 mM EDTA, 20 mM Tris-HCl, pH 8.1, 500 mM NaCl) and twice with TE-buffer (1 mM EDTA, 10 mM Tris-HCl, pH 8.1), chromatin was eluted (1% SDS, 0.1M NaHCO_3_)and recovered with Agencourt AMPure XP beads (Beckman Coulter). Bound chromatin and input DNA were treated with RNase H (New England Biolabs) and Proteinase K (Sigma-Aldrich) at 37 °C for 30 min. Immunoprecipitated chromatin was quantified by real-time PCR using the Maxima SYBR Green qPCR Master Mix (Thermo Scientific) and the ABI 7700 Sequence Detection System (Applied Biosystems). SDS 2.4 software (Applied Biosystems) was used for analysis. The specificity of each PCR reaction was confirmed by melting curve analysis using the Dissociation Curve software (Applied Biosystems). All chromatin immunoprecipitations and qPCRs were repeated at least three times, and representative results are shown. Primer sequences were HIV LTR Nuc0 forward: 5′ ATCTACCACACACAAGGCTAC 3′, HIV LTR Nuc0 reverse: 5′ GTACTAACTTGAAGCACCATCC 3′. HIV LTR Nuc1 forward: 5′ AGTGTGTGCCCGTCTGTTGT 3′, HIV LTR Nuc1 reverse: 5′ TTCGCTTTCAGGTCCCTGTT 3′.

### Statistical analysis

Two groups were compared using a two-tailed Student’s *t*-test. Multiple comparisons to a control were calculated using two-way ANOVA, followed by Dunnett’s test. Data are presented as mean ± SD for biological replicates. Statistical significance is indicated in all figures as follows, p-value: *p<0.05, **p<0.01, ***p<0.001, ****p<0.0001.

---

**Table S1.**
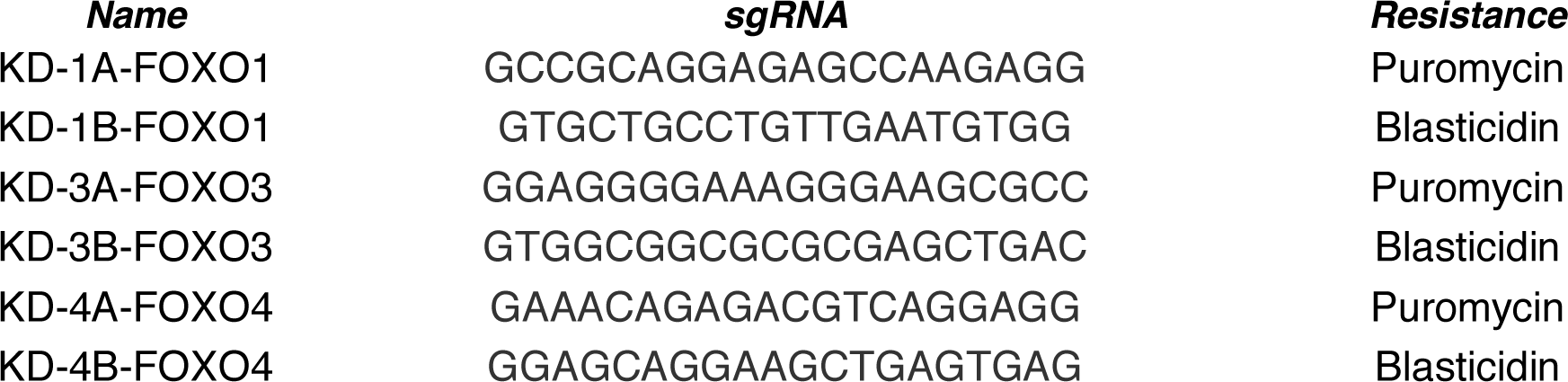
sgRNA list.

**Table S1.**
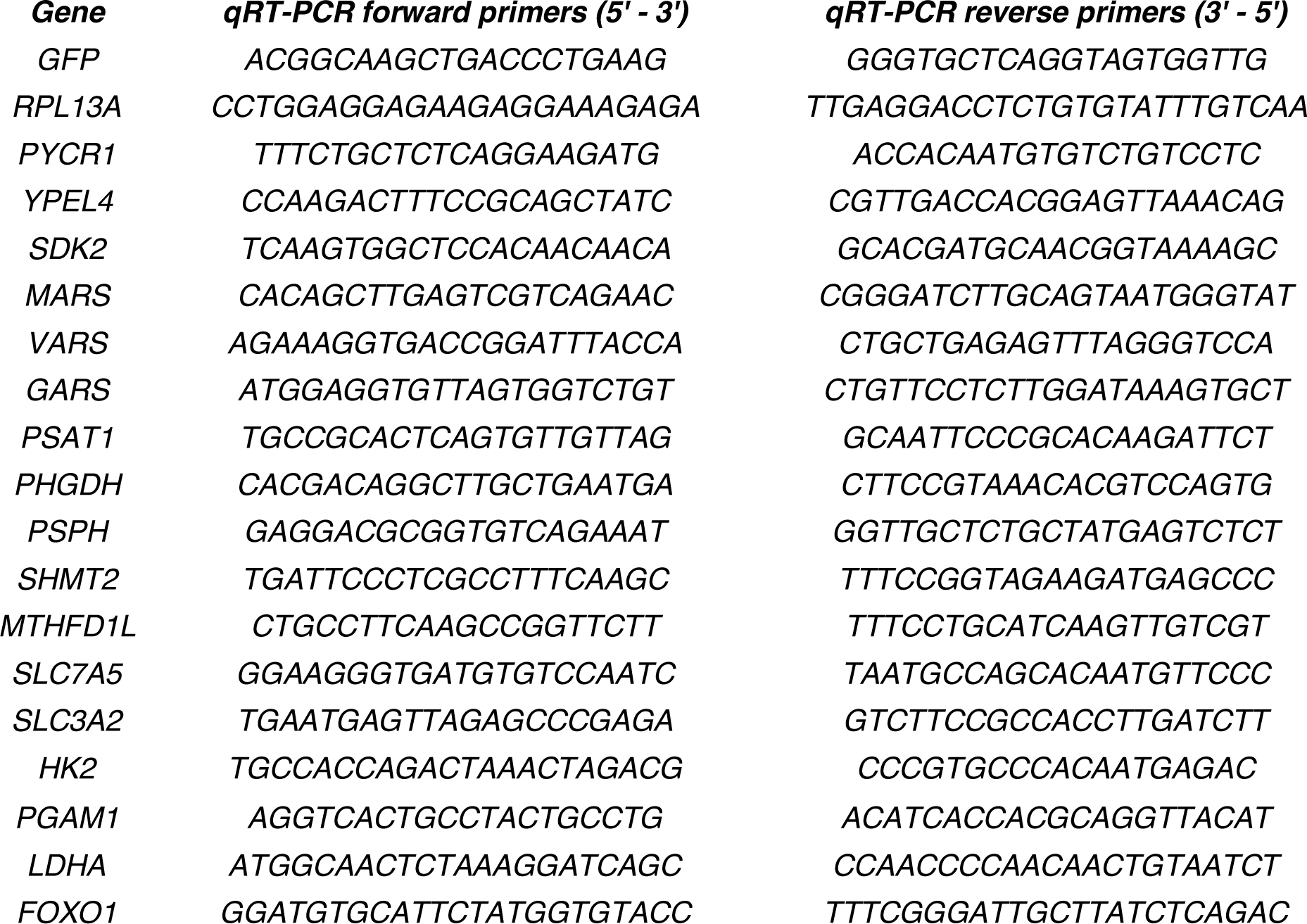
qRT-PCR primer list.

## Data availability

The complete array datasets of the RNA sequencing data associated with Figure 4a, b and Extended Data Figure 4 are available at the Gene Expression Omnibus with the accession number GSE129522.

## Acknowledgements

We thank all members of the Ott, Verdin and Pillai laboratories for helpful discussions, reagents and expertise. We thank, Dante Lacuadra and Austin Patel for assistance, Marielle Cavrois, Nandhini Raman, and the Gladstone Flow Cytometry Core for assistance with FACS, Natasha Carli, Jim McGuire and the Gladstone Genomics Core for assistance with RNA-Seq, John Carroll for graphics, Kathryn Claiborn and Brett Mensh for editorial assistance, and Veronica Fonseca and Lauren Weiser for administrative assistance. This work was supported by NIH/NIAID R01939 and NIH/NIAID R01985 to M.O., NIH/NIDA R01DA041742 and NIH/NIAID R01AI117864 to E.V., NIH/NIGMS R01GM117901 to S.K.P., and NIH P30 AI027763 to Flow Cytometry Core. Research was also supported as part of the amfAR Institute for HIV Cure Research, with funding from amfAR grant number 109301. D.B. was also funded by the Gilead HIV Cure Mentored Scientist Award from the amfAR Institute for HIV Cure Research at the UCSF AIDS Research Institute (ARI).

## Author contributions

A.V.-G. and M.O. designed and guided the study. A.V.-G., I.C. and R.P. designed, performed and analyzed most biochemical experiments. E.B. helped designing the project. D.B. performed ChIP assays. E.B. helped with primary cell experiments. T.T. performed CETSA assays. K.K., K.R., P.A.H., M.W., I.H., and A.C. performed some biochemical experiments. S.D. provided resources, and S.P., E.V. and M.O. provided the use of their laboratories and helped with data interpretation. The manuscript was written by A.V.-G. and M.O.

## Extended Figures

**Extended Data Figure 1.**
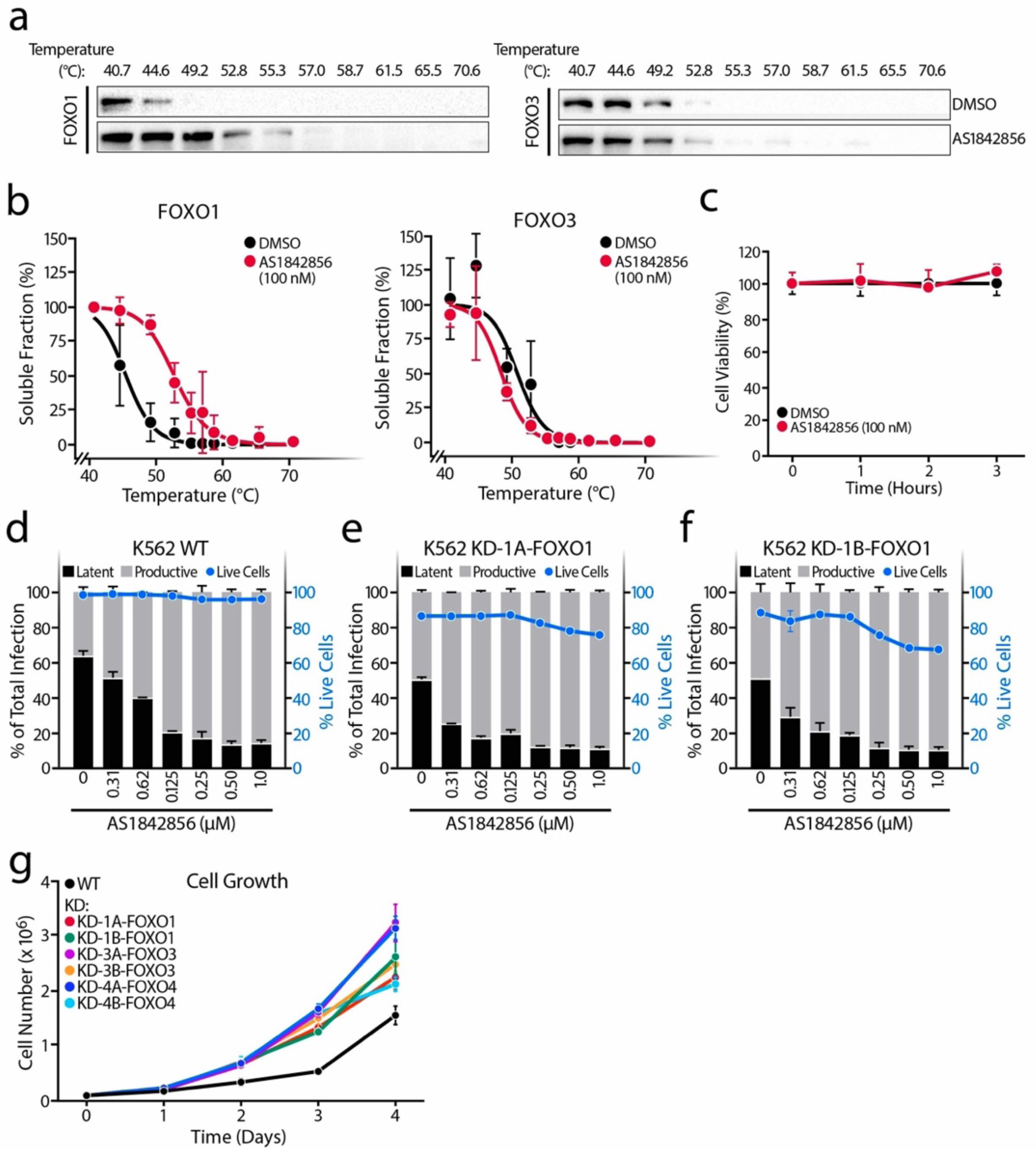
**a**, Representative western blots of CETSA assays in FOXO1 and FOXO3 in the presence or absence of 100 nM AS1842856. (b) FOXO1 and FOXO3 CETSA-melting curves upon the presence or absence of AS1842856 100 nM in K562 cells. Band intensities obtained from western blot analysis were normalized to the highest western blot signal which has been set to 100 %. Relative FOXO-band intensities were plotted against corresponding incubation temperatures and a nonlinear least-squares regression fit was applied. Data represent the mean ± SD of three individual experiments. **c**, Cell viability of CETSA assay in FOXO1 experiments assessed by Trypan Blue exclusion. **d-f**, Percentage of productive or latent cells relative to the total infection rate and cell viability upon increasing concentrations of AS1842856 treatment in the WT K562 (**d**) or in the FOXO1-knockdown cell lines KD-1A (**e**) and KD-1B (**f**). Data are represented as mean ± SD of two independent experiments. **g**, Cell growth analysis of WT, FOXO1, FOXO3 and FOXO4 knockdowns K562 cell lines.

**Extended Data Figure 2.**
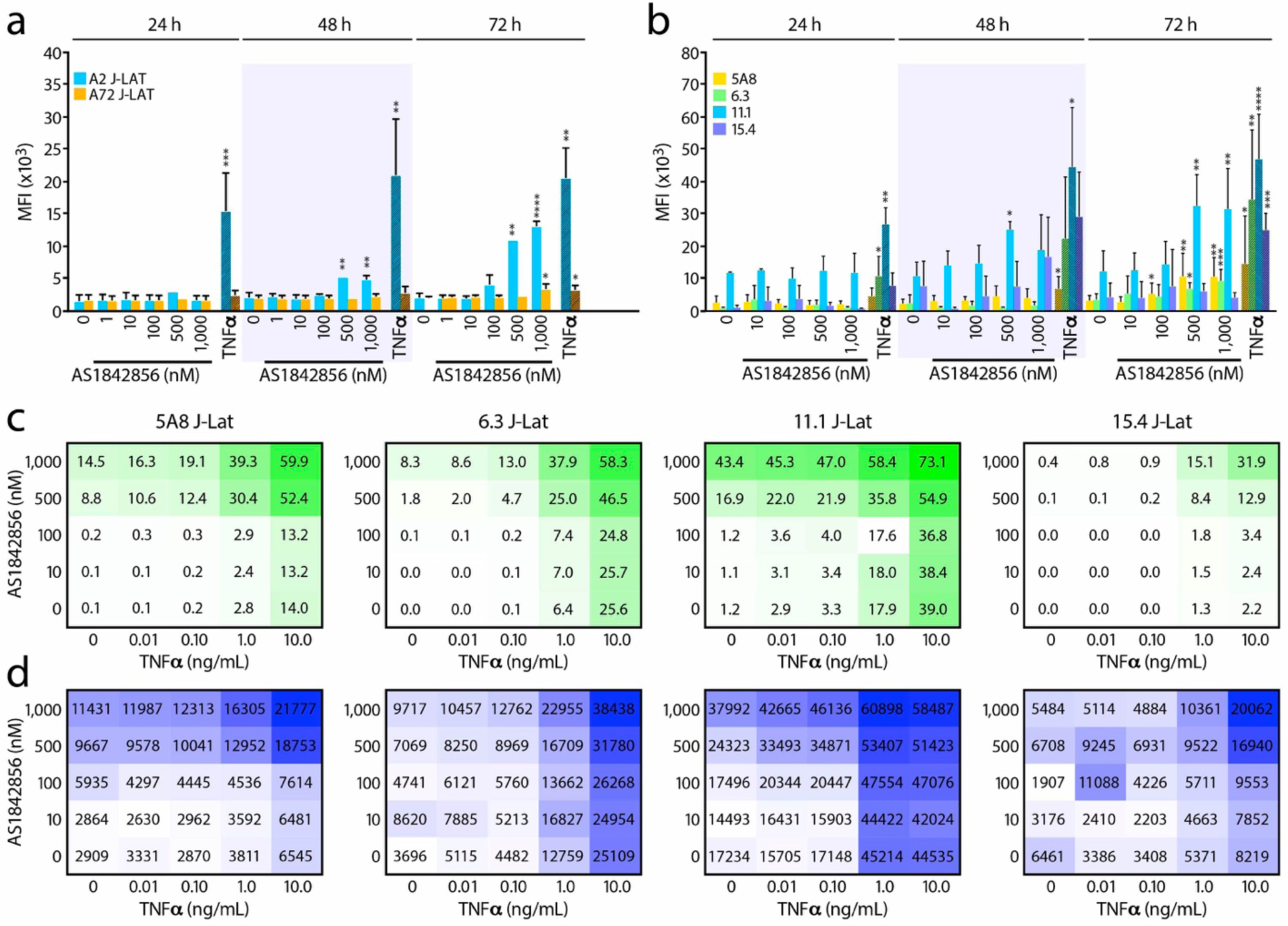
**a**, J-Lat cell lines A2 and A72 were treated with increasing concentrations of AS1842856 for 24, 48 and 72 h and HIV-GFP reactivation was analyzed by flow cytometry. HIV-GFP reactivation is reported as a Mean Intensity Fluorescence (MFI) of GFP-expressing cells. Data represent mean ± SD of at least three independent experiments. **b**, Same experiments as in A but performed in J-Lat cell lines 5A8, 6.3, 11.1 and 15.4. Data represent average ± SD of at least three independent experiments. 10 ng/mL TNFα was used as control. **c**, J-Lat cell lines 5A8, 6.3, 11.1 and 15.4 were treated for 72 h with increasing concentrations of both AS1842856 (Y-axis) and TNFα (X-axis) alone or in combination and analyzed by FACS. HIV-GFP reactivation is reported as a percentage of GFP-expressing cells (% GFP+ cells) or **d**, Mean Intensity Fluorescence (MFI). Data represent mean of three independent experiments.

**Extended Data Figure 3.**
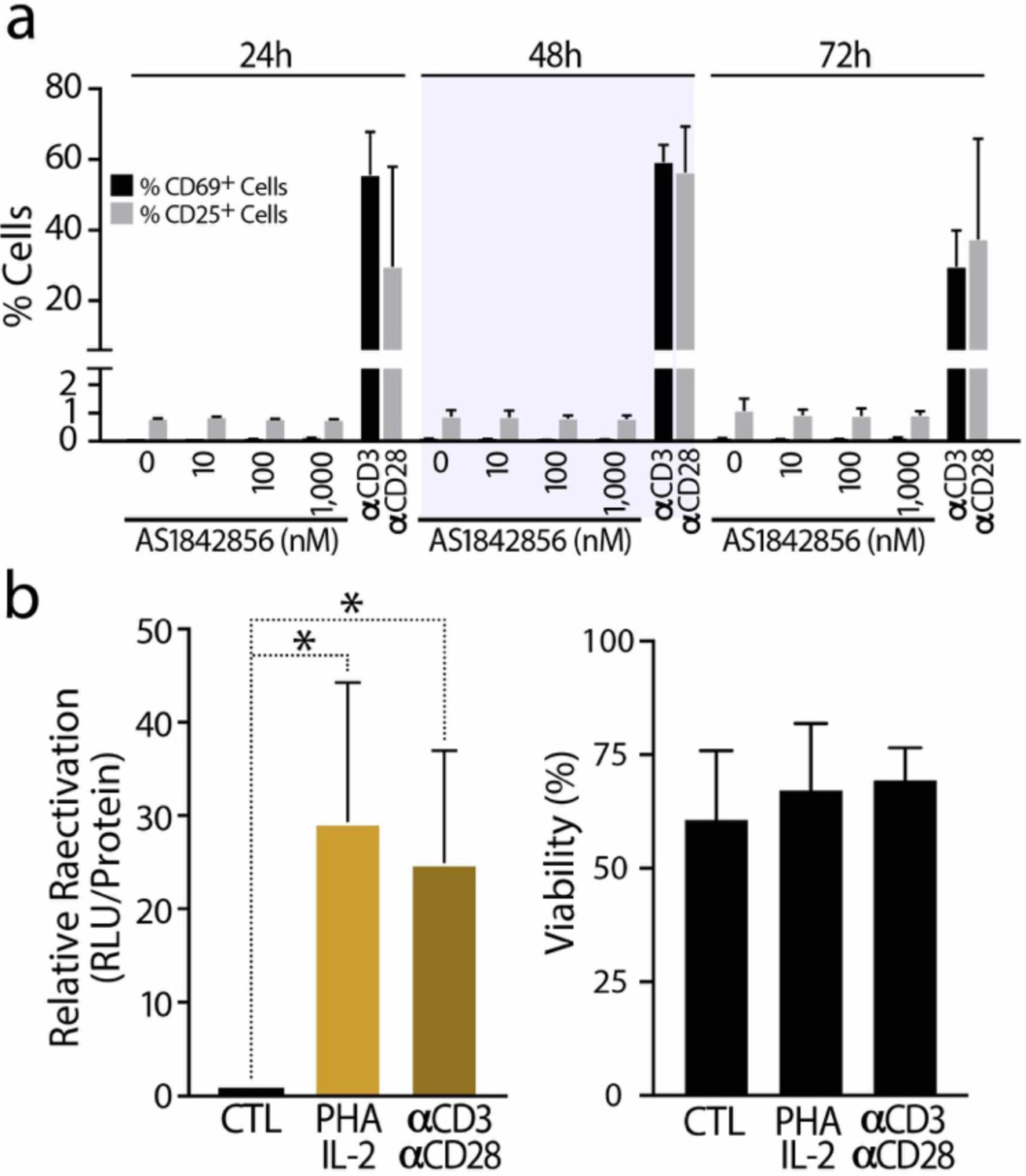
**a**, The cell surface CD69 and CD25 T cell activation markers were measured by FACS in CD4^+^ T cells upon AS1842856 treatment for 24, 48 and 72 h. 10 µg/mL αCD3 and 1 µg/mL αCD28 was used as control. Data is shown as mean of percentage of positive cells and as mean ± SD of two biological replicates. **b**, HIV reactivation was measured by luciferase activity and cell viability by flow cytometry assessed in CD4^+^ T cells purified from blood of healthy donors and infected with HIV_NL4-3 Luciferase_ and letting them rest for 6 days before reactivation was induced with 10 µg/mL PHA + 100 U/mL IL-2 and 10 µg/mL αCD3 + 1 µg/mL αCD28 for 72 h, in the presence of raltegravir (30 μM). Data represent average ± SD of at least nine independent experiments.

**Extended Data Figure 4.**
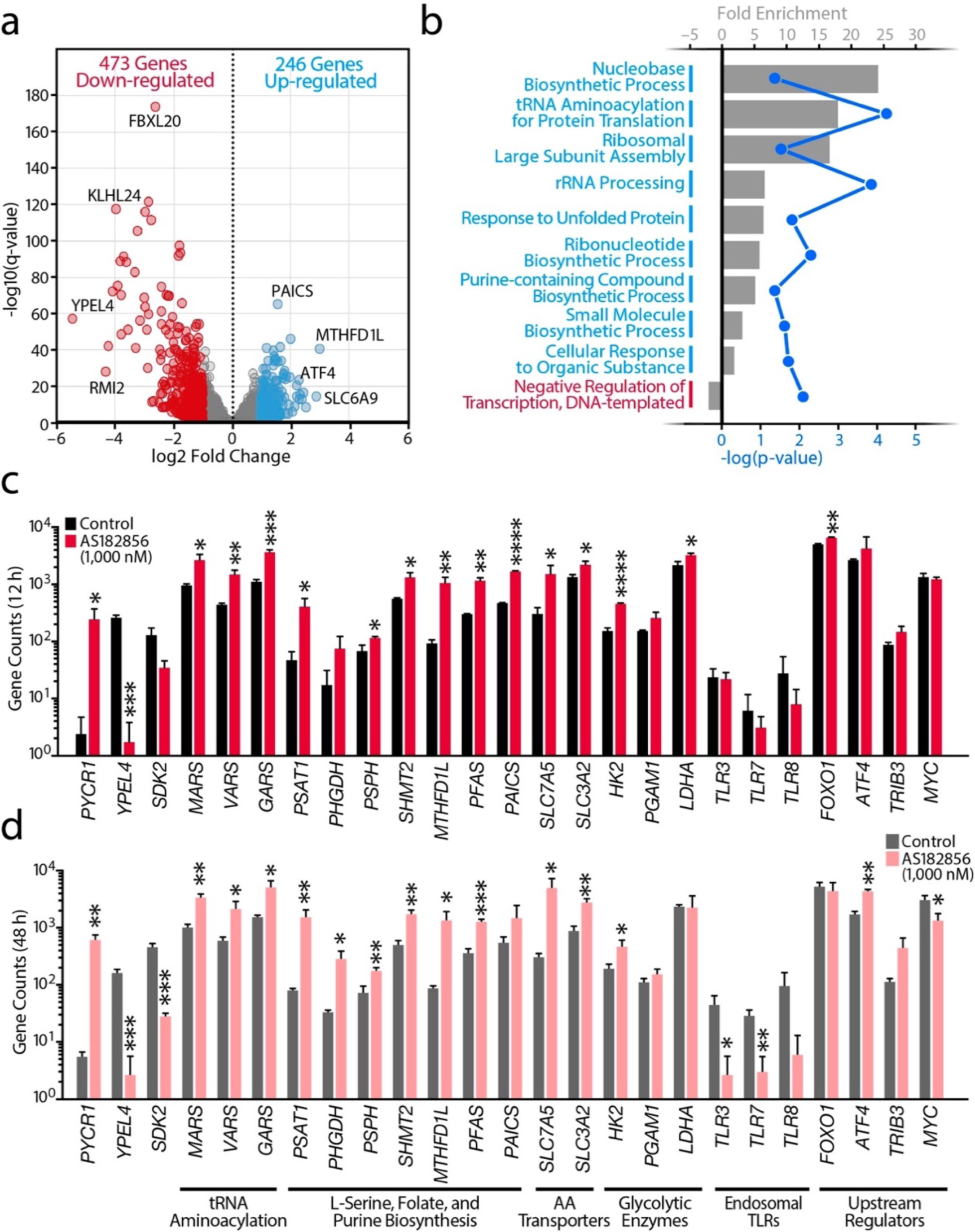
**a**, Volcano plot from the RNA-Seq data comparingCD4^+^ T cells treated with AS1842856 (1,000 nM) versus DMSO control for 12 h. Up-regulated (blue) or down-regulated (red) genes are q-value < 0.05 and log_2_ fold change ≥ 1 or ≤ −1, respectively. **b**, Analysis of the most dysregulated canonical pathways for the up- and down-regulated genes at 12 h. **c-d**, Confirmation of specific gene expression changes by mean gene counts from sequencing at 12 h (**c**) and 48 h (**d**). Data are represented as mean ± SD of three independent experiments.

**Extended Data Figure 5.**
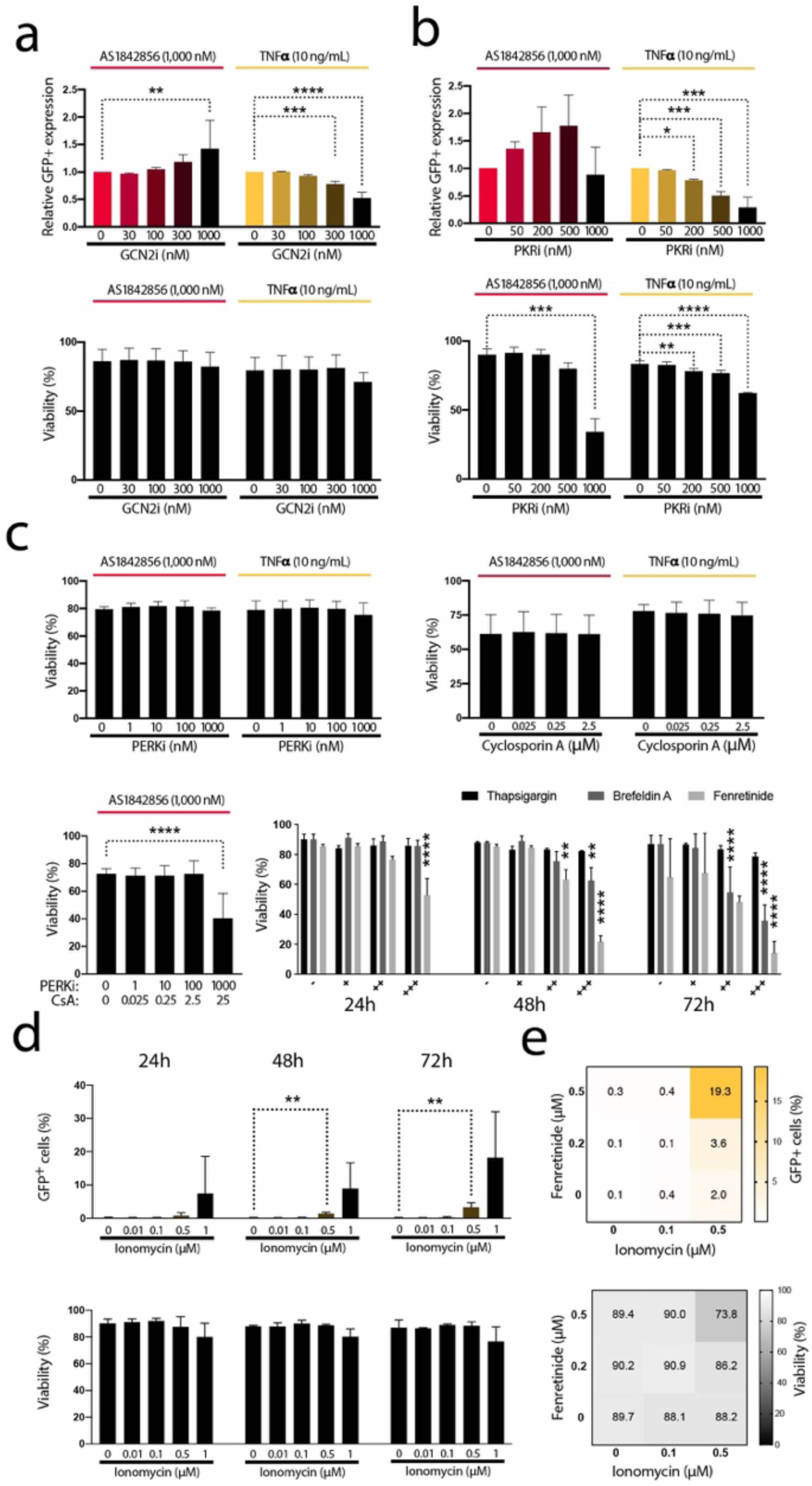
**a**, J-Lat cell line A58 was treated with increasing concentrations of GCN2i (A-92) in combination with 1,000 nM AS1842856 (72 h) or 10 ng/mL TNFα (24 h). In the upper panel, HIV-GFP reactivation was analyzed by FACS and relativized to the control. In the lower panel, histogram plots of percent live cells for each drug treatment are shown. **b**, Same experiment as in Extended Data. Fig.5a but treating cells with increasing concentrations of PKRi (Imidazolo-oxindole PKR inhibitor C16). Histogram plots representing viability are also shown. Data are represented by mean ± SD of three different donors. **c**, Histogram plots of percent live cells for each drug treatment are shown. Increasing concentrations of PERKi (GSK2656157 / PERK inhibitor II) (top left), Cyclosporin A (CsA) (top right), combined concentrations of PERKi and Cyclosporin A (bottom left) and increasing concentrations of Thapsigargin (0.01, 0.1, 1 µM), Brefeldin A (0.01, 0.1, 1 µg/mL) and Fenretinide (0.5, 2, 5 µM) for 24, 48 and 72 h (bottom right). Data represent mean ± SD of at least three independent experiments. **d**, J-Lat cell line A58 was treated with increasing concentrations of Ionomycin (0.01, 0.1, 0.5, 1 µM) for 24, 48 and 72 h and HIV-GFP reactivation (upper panel) and cell viability (lower panel) were analyzed by FACS. Data represent mean ± SD of at least three independent experiments. **e**, J-Lat cell line 5A8 was treated for 72 h with increasing concentrations of both Fenretinide (Y-axis) and Ionomycin (X-axis) alone or in combination and analyzed by FACS. HIV-GFP reactivation is reported as a percentage of GFP-expressing cells (% GFP+ cells) (upper panel) and viability was measured by FACS (bottom panel). Data represent average of three independent experiments.

